# OMA1 High-throughput Screen Reveals Protease Activation by Kinase Inhibitors

**DOI:** 10.1101/2021.10.02.462879

**Authors:** Marcel V. Alavi

## Abstract

Mitochondrial proteases are interesting but challenging drug targets for multifactorial diseases, such as neurodegeneration and cancer. The mitochondrial inner membrane protease OMA1 is a bona fide drug target for heart failure supported by data from human linkage analysis and animal disease models, but presumably relevant for more indications. OMA1 acts at the intersection of energy metabolism and stress signaling. The protease cleaves the structural protein OPA1, which organizes the cristae, as well as the signaling peptide DELE1, which can stimulate the integrated stress response. OMA1 shows little activity under physiological conditions but hydrolyzes OPA1 in mitochondria destined for mitophagy and during apoptosis. Little is known about OMA1, its structure has not been solved, let alone its context-dependent regulation. Autocatalytic processing and the lack of OMA1 inhibitors are thereby creating the biggest roadblocks. This study introduces a scalable, cellular OMA1 protease assay suitable for high-throughput drug screening. The assay utilizes an engineered luciferase targeted to the inner membrane as artificial OMA1 substrate, whereby the reporter signal inversely correlates to OMA1 activity. Testing different screening protocols and sampling different compound collections validated the reporter and demonstrated that both OMA1 activators as well as OMA1 inhibitors can be identified with the assay. Ten kinase-targeted cancer drugs triggered OMA1 in the assays, which suggests—considering cardiotoxicity as a rather common side-effect of this class of drugs—cross-reactivity with the OMA1 pathway.

## INTRODUCTION

The AAA proteases of the mitochondrial inner membrane are potential drug targets for indications across the disease spectrum. Human linkage analysis and data from a number of animal disease models validated the metalloendopeptidase OMA1 (from <u>o</u>verlapping function with <u>m</u>-A<u>A</u>A protease <u>1</u>) as *bona fide* drug target for heart failure.^1–8^ OMA1 is also an emerging oncotarget together with the i-AAA protease YME1L1.^9–11^ OMA1 partakes in mitochondrial quality control at the intersection of energy metabolism and stress response signaling. Among its substrates are the signaling molecule DELE1 and the cristae-shaping protein OPA1, which is also processed by the i-AAA protease.^12^

Mitochondria—often referred to as cellular powerhouses—can generate ATP by oxidative phosphorylation, whereby a proton gradient across the inner membrane powers ATP synthesis. The capacity of this electromotive force increases when the inner membranes align in orderly stacks, which are reinforced by the structural protein OPA1.^13–15^ This is illustrated by the parallel alignment of the cristae, for instance, in cardiomyocytes, which have been shown to rely on OPA1.^16–20^ These membrane stacks are rearranged—and OPA1 reorganized—in cells switching from the efficient generation of ATP to the resourceful synthesis of metabolites needed for cell growth and proliferation.^21, 22^ Examples for this include the fragmented appearance of mitochondria in activated immune cells, pluripotent stem cells and in cancer cells.^23–30^ It is still unclear, however, how the morphological changes of the inner membrane in response to energy-metabolic demands are regulated. It was suggested the i-AAA protease is downstream of AMPK.^31^ Another report suggested OMA1 is downstream of GSK3B.^32^

An additional layer of complexity is added by the mitochondrial quality control—a central mechanism for the maintenance of mitochondria health and homeostasis, which comprises balanced fission and fusion events. This dynamic behavior can sort out and separate damaged mitochondria, which are subsequently recycled by mitochondrial autophagy, aka mitophagy. DELE1 can thereby relay mitochondrial stress to the integrated stress response.^33, 34^ Severely harmed cells on the other hand are recycled in their entirety by initiating apoptotic cell death, which involves mitochondrial fragmentation along with OPA1 processing.^35–39^ The processes behind the judgement call between recycling of single mitochondria and entire cells are unknown, but could involve OMA1-dependent OPA1 cleavage, which is a signature event of both mitophagy and apoptosis. It is as yet unknown, however, how the activity of the OMA1 protease is regulated by the different environmental and cellular cues to initiate one program or the other.

The dilemma faced by OMA1 research is OMA1’s autocatalytic processing^40, 41^ and the lack of OMA1 inhibitors for the isolation of the protein. This creates a chicken-and-egg situation, because previously described OMA1 assays for the identification of OMA1 inhibitors relied on a short peptide coupled to a fluorescence resonance energy transfer (FRET) pair to measure protease activity. However, this peptide motive is recognized by many proteases and purified and functional OMA1 protease would be required for these assays to achieve OMA1 specificity. Yet, it is challenging to isolate functional OMA1 without (reversible) inhibitors. As a consequence, there are no OMA1 inhibitors available, which are urgently needed to advance OMA1 research, and which could potentially be useful starting points for a drug development program.

This study introduces the ‘Luke-S1’ reporter for cellular OMA1 protease activity assays. The Luke-S1 reporter is based on a split and permutated luciferase enzyme targeted to the mitochondrial inner membrane, where it represents an artificial OMA1 substrate. The Luke-S1 reporter establishes OMA1 specificity through its subcellular localization in the inner membrane. The signal thereby inversely correlates with OMA1 activity. Experimental OMA1 activation resulted in rapid Luke-S1 cleavage and elimination of its bioluminescence, which correlated with OPA1 hydrolysis in Hek293T cells stably expressing the reporter. Luke-S1 reporter cells produced a Z-prime score of 0.68 in 384-well format and thus are suitable for high-throughput drug screening. Testing different screening protocols and sampling different compound collections demonstrated that Luke-S1 can identify both OMA1 inhibitors as well as molecules that activate OMA1.

## RESULTS AND DISCUSSION

### OMA1-reporter design

All experiments of this study were conducted with Hek293T cells stably expressing the Luke-S1 reporter, which emits a signal inversely correlated with OMA1 activity (Figure 1). Luke-S1 is based on a modified NanoLuc complementation system.^42^ NanoLuc is an engineered luciferase, which can oxidize its cell-permeable substrate furimazine (8-benzyl-2-(furan-2-ylmethyl)-6-phenylimidazo[1,2-a]pyrazin-3-ol) thereby emitting light.^43^ The Luke-S1 reporter has the last 11 carboxy-terminal amino acids of NanoLuc (dubbed ‘SmBiT’) amino-terminally appended to the remaining 156 amino acids of the luciferase (‘LgBiT’) via a 24-amino acid linker encoding the human OPA1 *S1* cleavage site ‘T-A-F-R—A-T-D-R’ (P_4_-P_3_-P_2_-P_1_—P’_1_-P’_2_-P’_3_-P’_4_ with ‘—’ denoting the scissile bond; Figure 1A). Luke-S1 is targeted to the mitochondrial inner membrane by OPA1’s amino-terminus. The experiments were controlled by a reporter in which the *S1* site was replaced with a TEV site (‘E-N-L-Y-F-Q—S’; Luke-TEV) and by the native NanoLuc luciferase (herein referred to as ‘Luke’).

**Figure 1:**
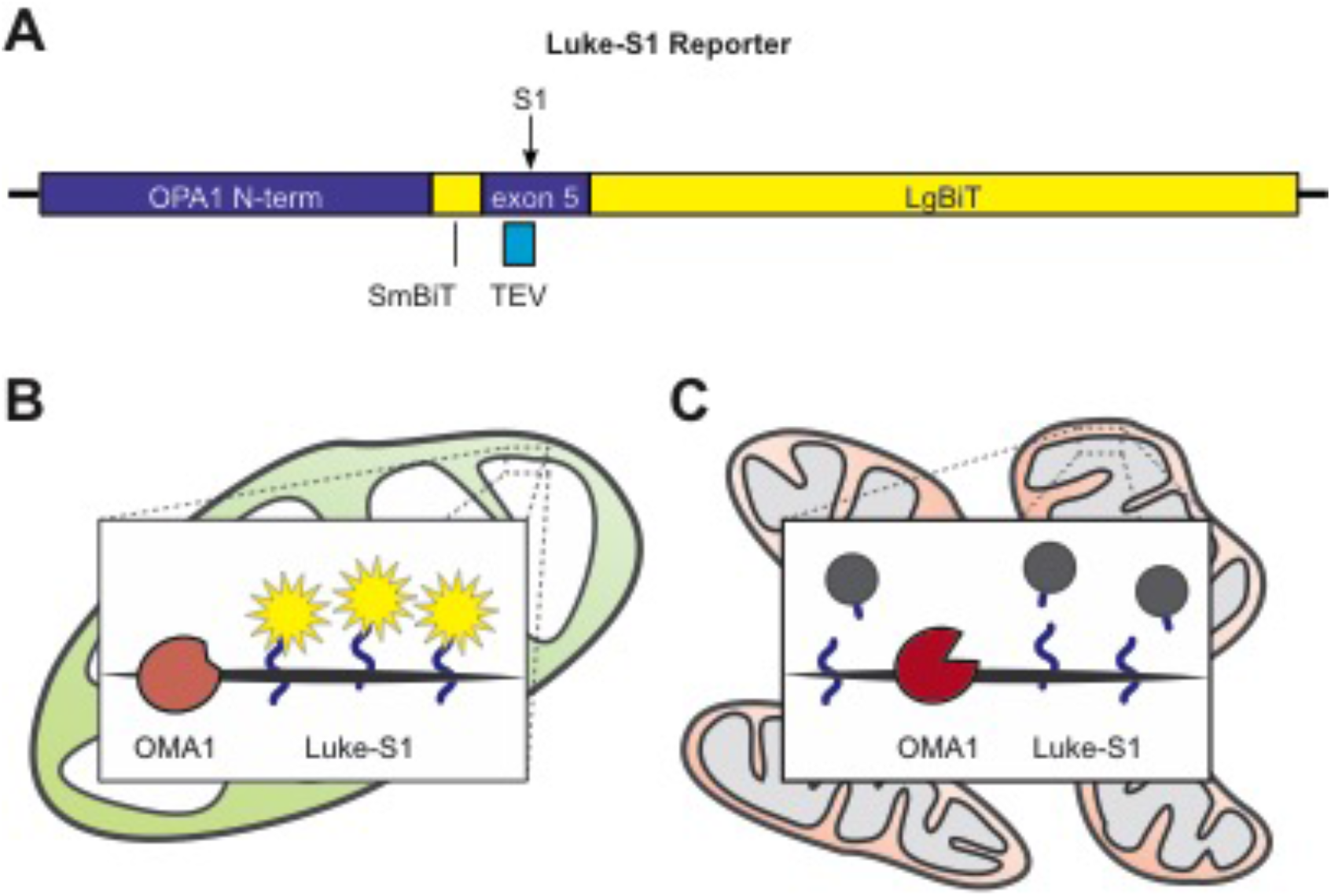
OMA1-reporter design and working principle. A reporter luciferase named Luke-S1 was engineered by combining portions of the OPA1 protein and the carboxy-terminus (SmBiT) and amino-terminus (LgBiT) of the NanoLuc luciferase to generate an artificial OMA1 substrate in the mitochondrial inner membrane (**A**). A TEV cleavage site replaces the *S1* cleavage site in the Luke-TEV reporter. Luke-S1’s bioluminescence inversely correlates with OMA1 activity. OMA1 is usually inactive and unhampered Luke-S1 can emit light (**B**). OMA1 becomes active in challenged cells and cleaves Luke-S1, which kills its bioluminescence (**C**).

### Context-dependent OMA1-reporter hydrolysis

OMA1 is usually inactive but becomes activated (and cleaves OPA1) in cells treated with the protonophore CCCP or the ionophore valinomycin.^44, 45^ Higher-weight L-OPA1 isoforms disappeared in Western blots of Hek293T cells after 30 minutes of treatment with 3 μM CCCP, which signified OMA1 activation (Figure 2A). Valinomycin is more potent and 100 nM for 30 minutes sufficed for L-OPA1 processing (Figure 2B). 3 μM CCCP and 100 nM valinomycin also induced Luke-S1 hydrolysis in Hek293T cells stably expressing the reporter (Figure 2C). The LgBiT antibody recognized a protein in untreated Luke-S1 and Luke-TEV cells migrating just below the 34 kDa standard. The predicted size of the full-length Luke-S1 and Luke-TEV reporter is 31.8 kDa. This band leveled off in Luke-S1 cells treated with CCCP or valinomycin and a band migrating above the 15 kDa standard became much more prominent, which—according to its size of approximately 19 kDa—corresponds to the luciferase hydrolyzed at the *S1* site. A faint 19 kDa Luke-S1 band even in unchallenged cells suggested OMA1 could have some basal activity or that other proteases recognize the reporter as well. Though Luke-S1 does not carry OPA1’s *S2* cleavage site, which is recognized by the i-AAA protease.^46, 47^

**Figure 2:**
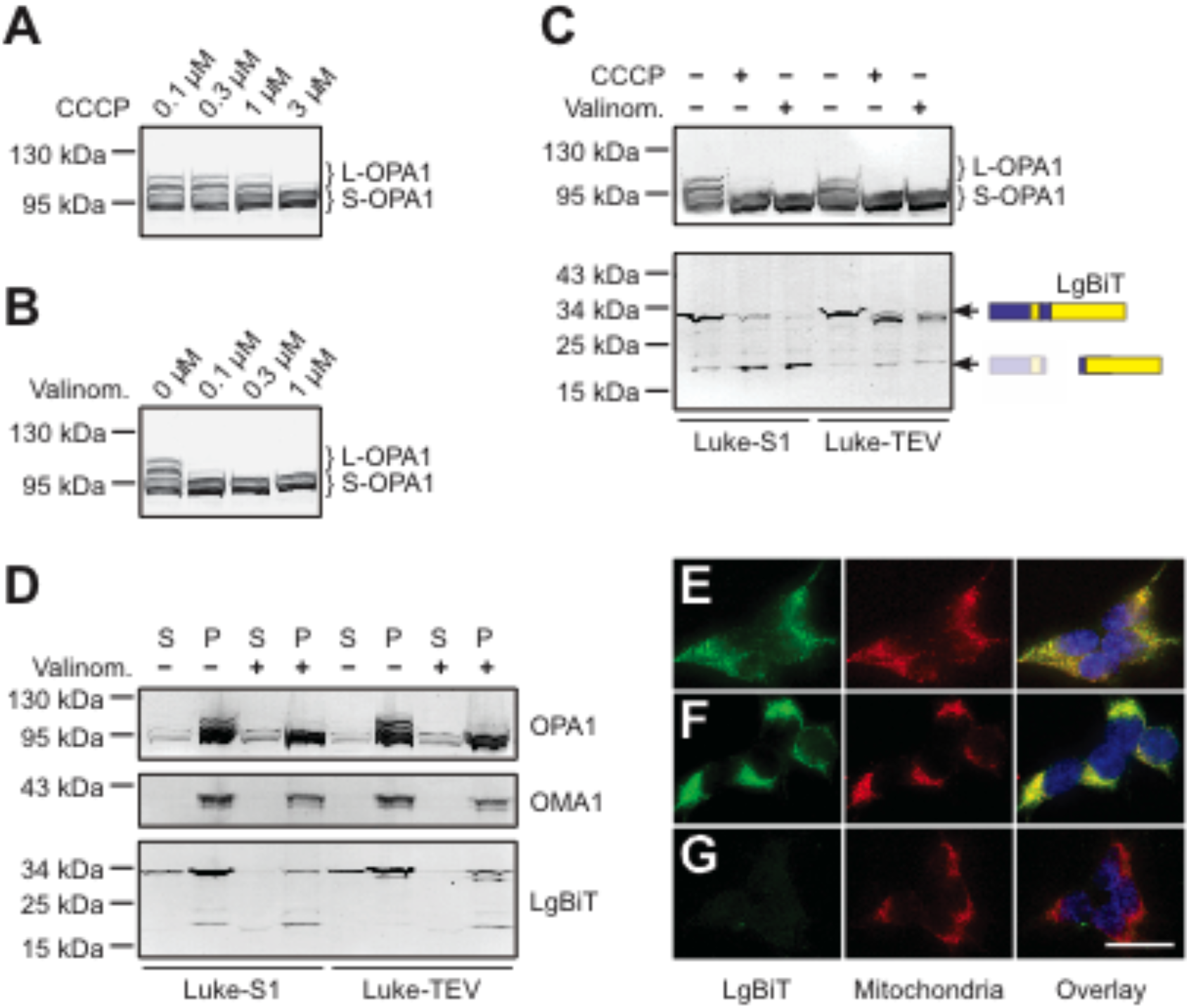
The Luke-S1 protein mirrors OPA1’s hydrolysis and subcellular localization. Western blots illustrate L-OPA1 proteolysis as proxy for OMA1 activity in Hek293T cells exposed to CCCP (**A**) or valinomycin (**B**). CCCP and valinomycin induced OPA1, Luke-S1 and Luke-TEV cleavage in Hek293T reporter cells (**C**). Cell fractionation by differential centrifugation located Luke-S1 and Luke-TEV together with OPA1 and OMA1 in the mitochondria-enriched pellet fraction (**D**; P, pellet; S, supernatant). Immunocytochemistry demonstrated LgBiT antibody co-labeling (green) with a mitochondrial dye (red) in Luke-S1 (**E**) and Luke-TEV cells (**F**), but not in Hek293T controls (**G**; bar: 25 μm).

Surprisingly, also Luke-TEV was cleaved in ionophore-treated cells. However, Luke-TEV showed a different cleavage pattern in Western blots with only a minor size reduction upon CCCP or valinomycin exposure (Figure 2C). Membrane proteases, such as the rhomboid protease, the gamma secretase or the signaling peptidase, are known to be rather promiscuous in their preference for cleavage sites and specificity is often established by gating mechanisms limiting access to the proteolytic cleft. This could also be the case for OMA1.^12^ For example, OMA1-expressing yeast strains could be characterized in vitro using FITC-casein as OMA1 substrate, which attests to OMA1’s promiscuity.^48, 49^ Another study described OMA1-dependent cleavage of misrouted PINK1 mutants^50^, which suggests that the placement of a protein in the inner membrane suffices for its recognition by the OMA1 protease. Cell fractionation by differential centrifugation found both Luke-S1 and Luke-TEV in mitochondria-enriched fractions together with OPA1 and OMA1 (Figure 2D). Also immunocytochemistry confirmed Luke-S1’s and Luke-TEV’s mitochondrial localization by co-labelling with a mitochondrial dye (Figure2E—G).

### OMA1-reporter specificity

Genetic ablation of the OMA1 protein could offset valinomycin’s effects on Luke-S1 reporter cells in Western blots and luciferase assays, verifying that OMA1 indeed processes Luke-S1. Valinomycin activated OMA1 in control siRNA-treated cells, which led to concomitant L-OPA1 and Luke-S1 hydrolysis (Figure 3A). As a result, L-OPA1 and Luke-S1 protein levels were significantly reduced to 18.6% ± 3.4 standard deviation (SD) and 7.2% ± 4.4 SD, respectively (Figures 3B & C; 2-way ANOVA: *p* < 0.0001). Interestingly, valinomycin also reduced OMA1 protein levels to 73.4% ± 6.7 SD, presumably due to its autocatalytic elimination.^40, 41^ Valinomycin’s effects on OPA1 and Luke-S1 were mitigated when OMA1 was depleted. siRNA-mediated knockdown reduced OMA1 protein levels to 31.9% ± 9.1 SD (Figure 3D). L-OPA1 levels were with 51.9% ± 9.9 SD significantly higher in valinomycin-treated OMA1 knockdown cells compared to control cells (Figure 3B). Also full-length Luke-S1 was more abundant in these samples (28.8% ± 9.1 SD), though without reaching statistical significance (Figure 3C). These results could be recapitulated in Luke-S1 luciferase assays. Valinomycin significantly reduced bioluminescence in Luke-S1 control cells but not in cells treated with OMA1 siRNA (Figure 3E). Previous studies reported that the chelator phenanthroline can inhibit the strictly zinc-dependent OMA1 protease.^46, 51, 52^ However, phenanthroline and other chelating drugs showed cytotoxicity in the Luke-S1 assays before a critical concentration for OMA1 inhibition could be reached (data not shown). Still, the OMA1 knockdown experiments confirmed that Luke-S1 is processed by OMA1.

**Figure 3:**
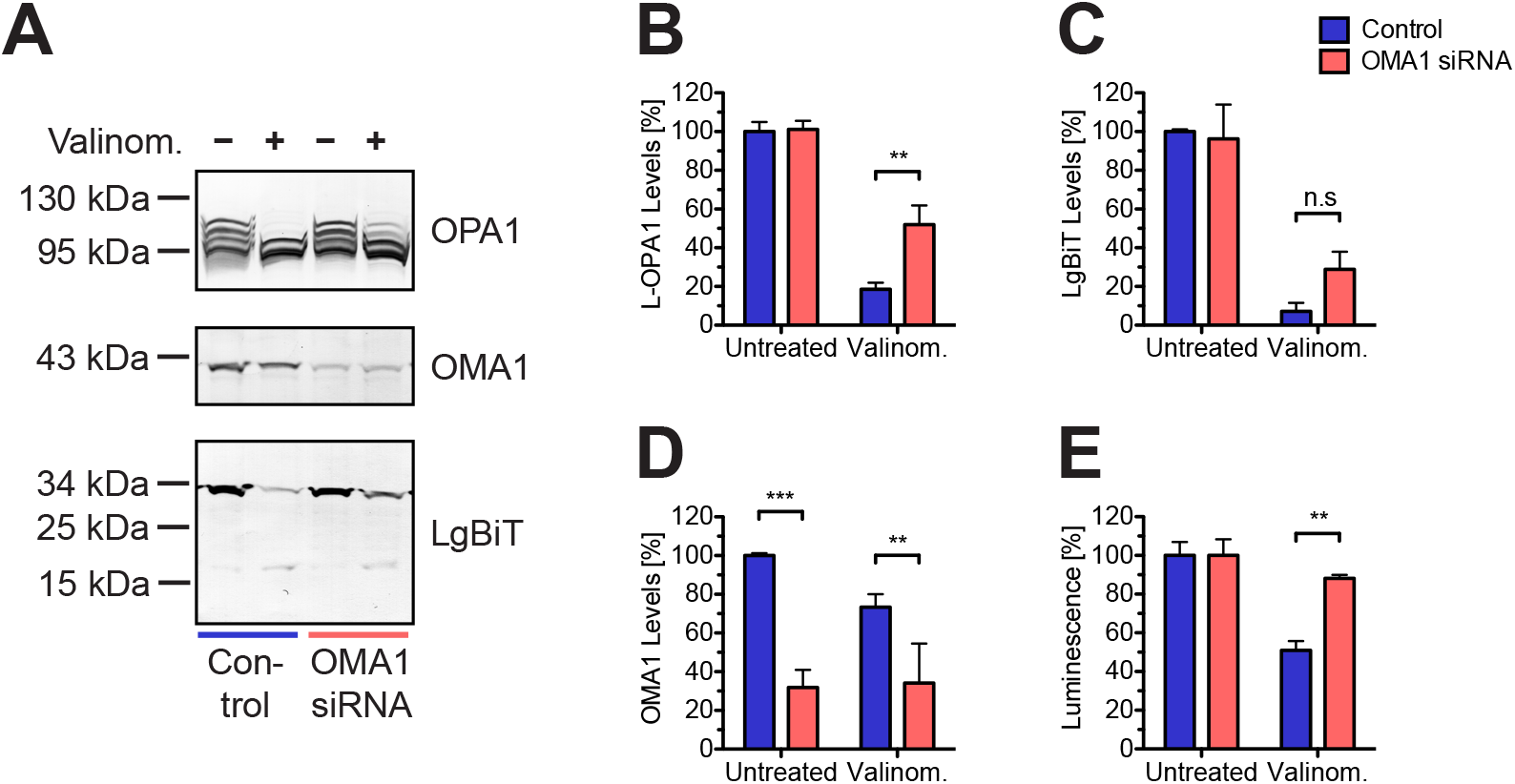
Genetic OMA1 ablation validates Luke-S1 reporter specificity. siRNA mediated OMA1 knockdown prevented valinomycin-induced OPA1 and Luke-S1 processing in Western blots (**A**) and diminished its effects in luciferase assays. Densitometric quantification demonstrated that significantly higher L-OPA1 protein levels in valinomycin-treated OMA1 knockdown cells (**B**) corresponded to higher LgBiT levels in these samples (**C**). OMA1 protein levels were thereby reduced by about 70% in the knockout samples (**D**). Valinomycin reduced the signal in Luke-S1 luciferase assays, but not when OMA1 was depleted in these cells (**E**). Bar graphs show mean ± SD; n = 3; 2-way ANOVA: *p* > 0.05 (n.s.); *p* < 0.001 (**); *p* < 0.0001 (***).

### OMA1-reporter characterization

The permutated reporter assembled into a functional luciferase when stably expressed in Hek293T cells. The basic characterization of the reporter enzymes in luciferase assays with increasing luciferase substrate concentrations revealed Michaelis-Menten substrate affinities (K_M_) of 37.9 μM ± 7.2 SD for Luke-S1 and 30.1 μM ± 6.8 SD for Luke-TEV. These were in the range of the K_M_ of the native NanoLuc enzyme (K_M_[Luke]: 31.3 μM ± 5.2 SD; Figure 4A—C). However, V_max_ was notably reduced compared to Luke by about an order of magnitude (V_max_[Luke-S1]: 119.7 ± 8.9 SD; V_max_[Luke-TEV]: 52.9 ± 4.2 SD; V_max_[Luke]: 536.8 ± 33.1 SD).

**Figure 4:**
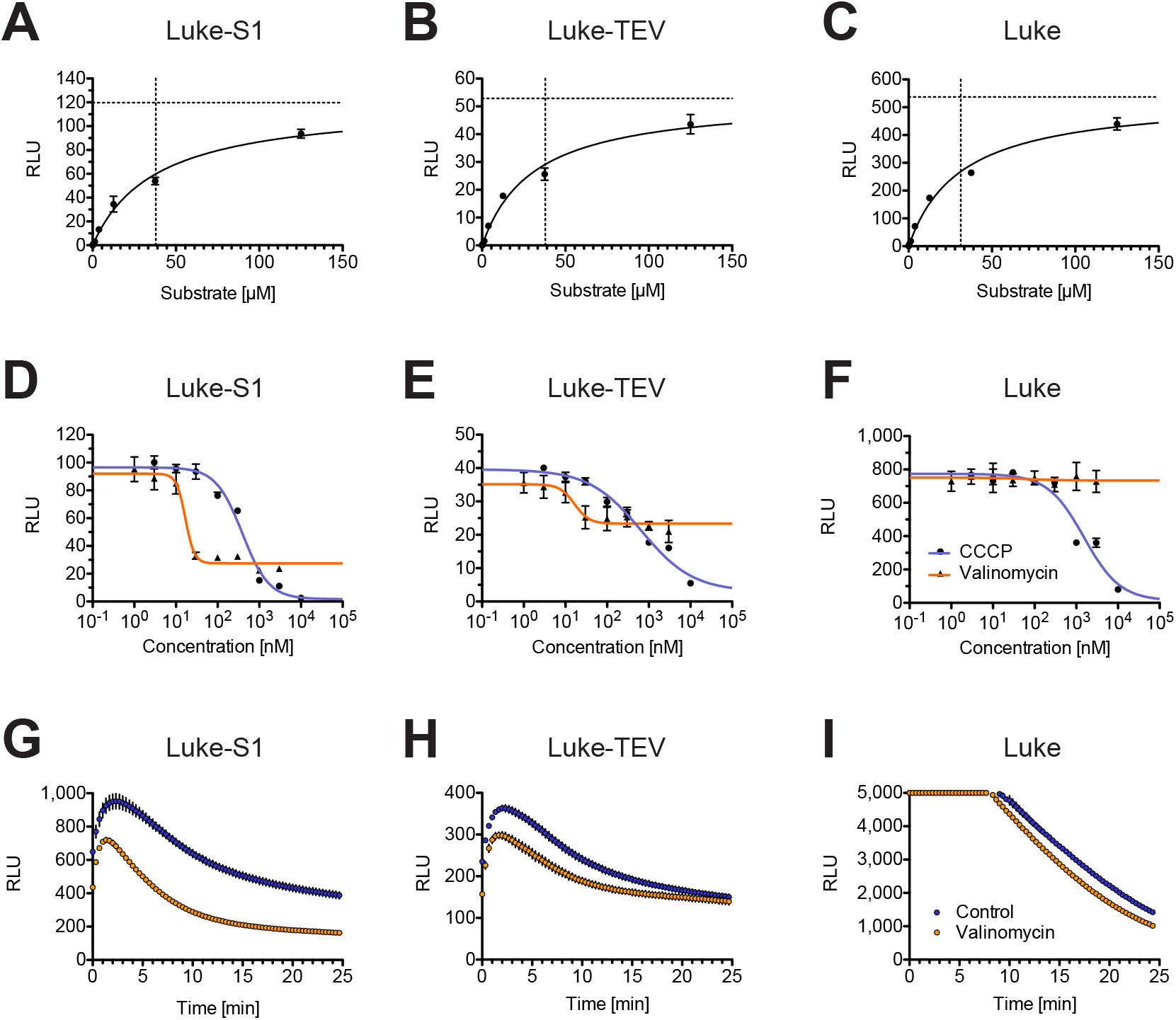
Characterization of the OMA1-reporter in luciferase assays. Plotting Michaelis-Menten enzyme kinetics revealed comparable K_M_ values for all 3 reporters, but lower V_max_ for Luke-S1 (**A**) and Luke-TEV (**B**) compared to the native luciferase Luke (**C**). Valinomycin reduced in a dose-dependent manner Luke-S1’s bioluminescence (**D**) and to a lesser extend Luke-TEV’s signal (**E**), without showing an effect on Luke control cells (**F**). CCCP obliterated the bioluminescence of all 3 reporter, including Luke, most likely by interfering with the luciferase (**D—F**). The dynamic behavior of Luke-S1 (**G**) and Luke-TEV (**H**) simultaneously exposed to valinomycin and luciferase substrate was rapid with differences being apparent from the beginning of the recordings. Valinomycin again had no effect on Luke in these time-course experiments (**I**). Note, signal decay in this assay is a multipart effect of reporter enzyme inactivation and substrate depletion.

Luciferase assays with CCCP or valinomycin presented a dose-dependent signal reduction, whereby CCCP also reduced bioluminescence in Luke controls. CCCP’s half maximal effective concentration (EC_50_) in Luke-S1 assays was 390.8 nM (95% confidence interval: 281—542 nM; *R^2^*=0.98) and valinomycin had an EC_50_ of 16.8 nM (95% confidence interval: 12.1—23.4 nM; *R^2^*=0.97; Figure 4D). CCCP also reduced the signal produced by Luke-TEV with an EC_50_ of 653.0 nM (95% confidence interval: 330—1293 nM; *R^2^*=0.97), while similar valinomycin concentrations that extinguished Luke-S1’s signal produced only a minor signal reduction of about 30% in Luke-TEV cells (EC_50_: 16.5 nM; 95% confidence interval: 7.4—36.6 nM; *R^2^*=0.81; Figure 4E). Valinomycin had no effect in Luke assays, but CCCP eliminated Luke’s bioluminescence (EC_50_: 1.5 μM; 95% confidence interval: 0.7—3.3 μM; *R^2^*=0.93; Figures 4F), which suggests that CCCP can engage the luciferase. An unrelated study noted previously CCCP’s assay interference with their NanoLuc-based reporter systems^53^, which affirms CCCP’s alleged cross reactivity with the luciferase. Nevertheless, valinomycin did not show such interference even in the micromolar range. For this reason, valinomycin at a concentration about five-times its EC_50_ (i.e. 100 nM) was considered a good trigger stimulus for all experiments.

To better understand the dynamic nature of the OMA1 assay, reporter cells were exposed to luciferase substrate without or with valinomycin and bioluminescence was recorded immediately in real-time in 30-second intervals (Figure 4G—I). Luke-S1 showed a marked signal reduction from the start of the measurement compared to cells without valinomycin. (It required about two minutes from adding the substrate, mixing and spinning the plate to start the measurement.) The signal also declined more rapidly during the first 15 minutes in the presence of valinomycin before the signal stabilized at a significantly lower intensity of about 40% (Figure 4G). Luke-TEV on the other hand showed a much smaller difference from the beginning of the measurement and after 15 minutes there were no differences between cells without and with valinomycin anymore (Figure 4H). The strong signal intensity of Luke saturated the photomultiplier tube of the instrument in the beginning (Figure 4I). Nonetheless, after 10 minutes there were no notable differences in bioluminescence levels or signal decay between the two treatment conditions. Luke’s signal decay was thereby most likely caused by substrate depletion due to NanoLuc’s much higher V_max_. Follow-up studies indicated signal recovery of Luke-S1 reporter cells two days after OMA1 activation (data not shown). Even so, additional experiments would be needed to clarify how cell proliferation, restoration of OMA1, and Luke-S1 renewal are connected. Overall, 30 minutes of 100 nM valinomycin provided enough time for OMA1 activation, reporter cleavage and stabilization of the signal.

### OMA1-reporter pilot screens

To further evaluate the Luke-S1 reporter, different screening protocols were implemented and diverse compound libraries sampled. A first screen had the goal to gauge how many molecules would significantly diminish Luke-S1’s bioluminescence (Figure 5A). 3,500 chemically diverse compounds were sampled in 384-well plate format by incubating Luke-S1 cells for 60 minutes with 10 μM of each test compound. Luciferase substrate was then added and bioluminescence measured. All signals were normalized to untreated cells and any molecules that would reduce bioluminescence by more than 3 SD were considered hits. The bioluminescence of untreated cells in 352 wells across 11 plates followed a normal distribution; the average signal was 100% ± 9.35 SD and the 3 SD-significance threshold was set accordingly at 72.0% (Figure 5A, dotted line). 11 of the 3,500 molecules (0.3%) significantly reduced Luke-S1’s bioluminescence in this screen.

**Figure 5:**
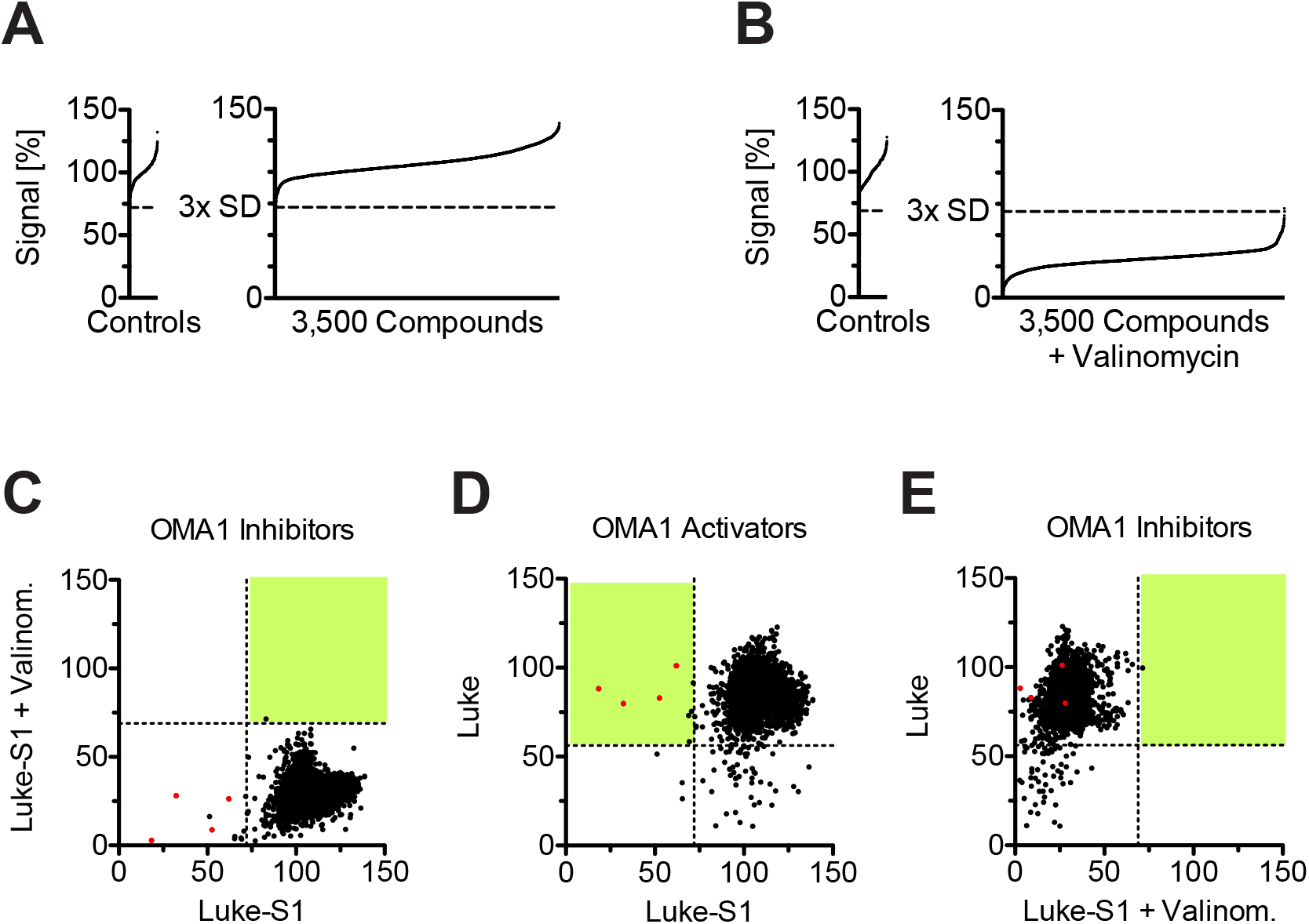
Proof of principle of OMA1-reporter assays. Luke-S1 cells were exposed to 3,500 compounds in search of molecules that would lower the reporter signal by at least 3 SD of untreated controls (**A**) or that would maintain the reporter signal within the 3 SD upon valinomycin exposure (**B**). Plotting the data of both screens in 2 dimensions would place OMA1 inhibitors in the upper right quadrant (**C**). The 3,500 compounds were also tested on Luke cells. OMA1 activators, i.e. molecules that reduced the signal of Luke-S1 but not Luke controls, moved to the upper left quadrant of the 2D scatterplot (**D**, red dots). All hits from the Luke control screen had no influence on valinomycin-treated Luke-S1 cells and hence moved into the lower left quadrant (**E**).

An independent screen searched for molecules that would counteract valinomycin-induced OMA1 activation and hence sustain bioluminescence upon valinomycin exposure (Figure 5B). Luke-S1 cells were again incubated with the 3,500 test chemicals for 60 minutes, after which valinomycin was added for another 30 minutes before bioluminescence was recorded. All signals were normalized to untreated controls (100% ± 10.36 SD) and the significance level was defined as 68.9% (Figure 5B, dotted line). The average signal of all 3,500 valinomycin-treated samples was 30.7% ± 7.04 SD with one molecule marginally achieving significance. Plotting the data of both screens in two dimensions placed potential OMA1 activators in the lower left quadrant and potential OMA1 inhibitors in the top right quadrant (Figure 5C).

Because the Luke-S1 assay would also pick up any chemicals that interfere with the luciferase (e.g., CCCP), the 3,500 molecules were also tested on Luke control cells. To this end, Luke cells were treated like the Luke-S1 cells with 10 μM compounds for 60 minutes. The data obtained with Luke cells was noisier due to Luke’s much higher V_max_, which caused considerable plate drift (see Supplementary Information, Figure S1). The average signal of untreated Luke cells was 100% ± 14.62 SD clustering in two peaks corresponding to the untreated controls in column #2 and #23 of each plate, respectively. The 3 SD-hit threshold was set at 56.2%. 56 of the 3,500 molecules (1.6%) crossed this threshold and significantly reduced Luke’s bioluminescence. OMA1 inhibitors could be stratified from molecules that suppressed the luciferase by plotting Luke-S1 against Luke in 2D scatterplots (Figure 5D, top left quadrant). All molecules that significantly reduced Luke’s bioluminescence still responded to valinomycin in Luke-S1 cells. That is, they moved from the bottom right quadrant in Figure 5D to the bottom left quadrant in Figure 5E. For this reason, the number of molecules that interfered with the luciferase might actually be much lower than the 1.6% and presumably even as low as 0.09% (i.e. the 3 molecules in the bottom left quadrant of Figure 5D). This result is at odds with an independent study, which reported that 2.7% of their 42,000 sampled chemicals inhibited NanoLuc by at least 30%.^54^ It is possible that some of the 3,500 molecules were already degraded. The library was obtained from the NCI, where it had been stored for more than 10 years. Still, these pilot studies validated the OMA1-reporter and provided proof of concept that it is feasible to identify OMA1 activators and OMA1 inhibitors with the Luke-S1 reporter.

### OMA1 inhibitor screen

To identify OMA1 inhibitors an additional twenty-thousand chemicals were screened. Again, Luke-S1 cells were preincubated with test molecules for 60 minutes before valinomycin was added for another 30 to 60 minutes. Bioluminescence was recorded thereafter and all measurements were normalized to untreated cells. The average of 928 controls across 58 plates was 100% ± 10.4 SD and the hit threshold was set at 68.8% (Figure 6A, dotted line; Figure 6B illustrates the plate layout). The controls were not normally distributed but had a positive tail (Shapiro-Wilk normality test: *p* < 0.0001; Figure 6C), which was ascribed to higher signals in the top and bottom rows maybe caused by uneven temperature distribution across the plates. The average signal of the 20,416 test molecules after the addition of valinomycin was 29.6% ± 9.3 SD (Figure 6D). Six of the 20,416 test compounds (0.03%) passed the hit threshold with signals from 69.2% to 80.6% (Figure 6A). Cellular screening assays depend on cell permeability and are confounded by cytotoxicity, which increases stringency and which may explain the rather low hit rate of 0.03%.

**Figure 6:**
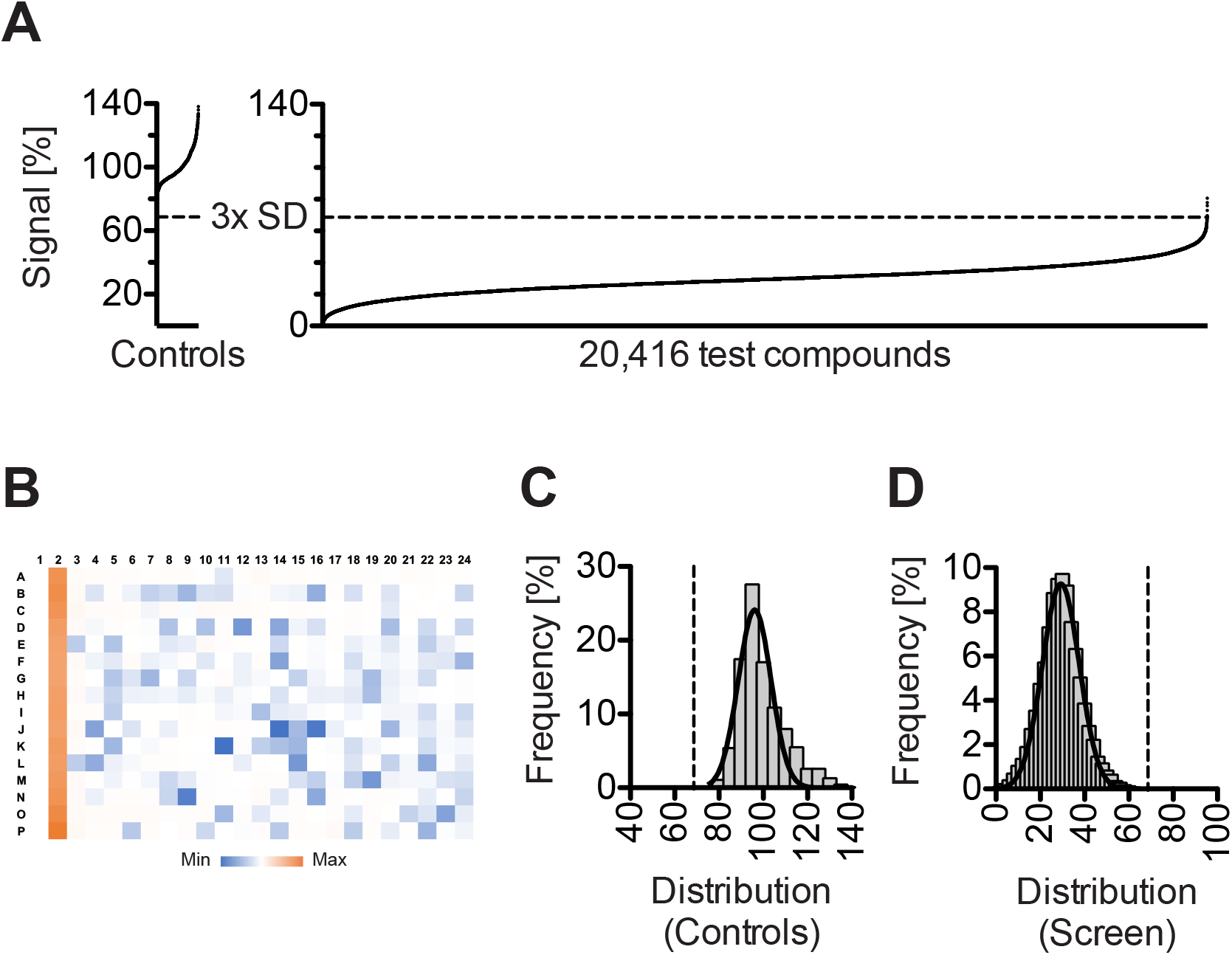
Screening for OMA1 inhibitors. 6 molecules sustained luminescence in valinomycin-treated Luke-S1 cells within 3 SD of untreated controls (A). A heatmap illustrates the plate layout with untreated controls in column #2 (B). The 928 controls did not follow a normal distribution but were skewed to the right (C), while the signal of the 20,416 followed a Gaussian distribution (D; *R^2^*=0.99).

### OMA1 activator screen

A number of cancer drugs, such as cisplatin and sorafenib, were reported to promote OMA1-dependent OPA1 cleavage^55, 56^, which provided the rational to test all 166 FDA-approved small molecule cancer drugs. To this end, Luke-S1 reporter cells were incubated with 10 μM of each drug for 60 minutes, after which luciferase substrate was added and bioluminescence measured. The signal intensity was normalized to untreated cells (100% ± 12.5 SD; valinomycin treated controls: 18.5% ± 5.6 SD) and the significance level was defined as a signal drop of ≥ 37.5 for this screen. 30 of the 166 approved cancer drugs (18.1%) reduced bioluminescence by 37.5% to 85.6%. 18 of the 30 hits were kinase inhibitors (60.0%), which constituted only 39.2% of all the drugs in this collection (65 of 166). This over representation was statistically significant (Fisher’s exact test: *p* = 0.013). Follow-up experiments with a 12-point titration curve could confirm OMA1 activation by 10 kinase inhibitors with EC_50_ values from 5 μM to 100 μM (Figure 7A—J). However, only the two most potent of these 10 drugs, Sorafenib (EC_50_: 4.8 μM; 95% confidence interval: 2.6—9.1 μM; *R^2^*=0.99) and Ceritinib (EC_50_: 10.6 μM; 95% confidence interval: 3.8—29.4 μM; *R^2^*=0.89), showed OPA1 and Luke-S1 processing in Western blots when cells were cultured for 3 hours with 50 μM of the drug (Figure 7K). A higher sensitivity of the Luke-S1 assay than probing Luke-S1 by Western blots was also noted for valinomycin. Why Luke-S1 assays showed a higher sensitivity than Western blots is unclear but might be attributed to the enzymatic signal amplification by the luciferase. The data for the other drugs were less conclusive (Supplementary Information, Figure S2). The antiestrogen tamoxifen is worth mentioning with an EC_50_ of 7.6 μM (95% confidence interval: 6.4—9.0 μM; *R^2^*=0.99). 2 drugs showed no effect in the Luke-S1 assay even at 100 μM. And the remaining 17 drugs had an EC_50_ > 100 μM or diminished also Luke’s bioluminescence so that no conclusions could be drawn. The anthracycline doxorubicin stood out from the latter because its 3 analogs daunorubicin, idarubicin and valrubicin also showed assay interference with EC_50_ values ranging from 8.0 μM to 19.0 μM (Supplementary Information, Figure S2).

**Figure 7:**
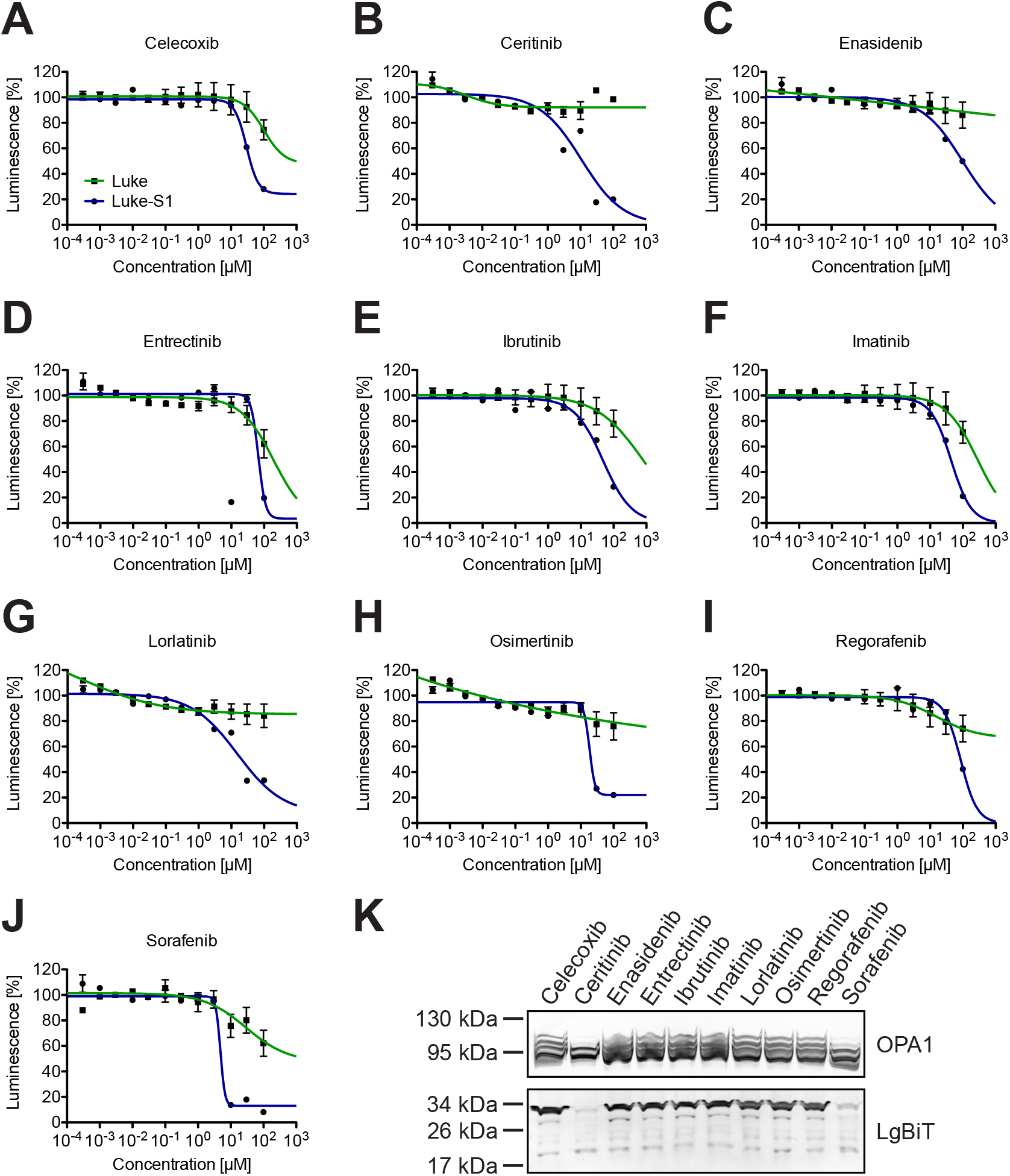
Luke-S1 picks up 10 kinase-targeted cancer drugs. The denoted cancer drugs triggered within an hour dose-dependent signal changes in the Luke-S1 assay (blue line), but showed only little assay interference in the Luke control assay (green line; **A—J**). Western blots revealed clearly discernable OPA1 and Luke-S1 processing (LgBiT) for the two most potent drugs after 3 hours of incubation (**K**).

## CONCLUSIONS

Luke-S1 is a suitable reporter to screen for both OMA1 inhibitors and OMA1 activators. An unexpected result was the number of kinase-targeted cancer drugs that showed up in the Luke-S1 assay. These kinase inhibitors engage various kinases to curb tumor growth and metastasis. It is not clear, however, whether the observed effects were a result of a direct interaction with OMA1 or the consequence of signaling events that just converged in OMA1 activation. The effect in Luke-S1 assays was rather rapid and already detectable after 60 minutes, which would speak for the former. However, just 2 of the 10 drugs led to distinct OPA1 and Luke-S1 cleavage in Western blots—and only after 3 hours of 50 μM, which would speak for the latter. AMPK and GSK3B could be potential mediators in such signaling events.^31, 32^ OPA1 has also been connected to NF-κB signaling.^57^ OMA1’s cancer connection has been recognized previously. For example, cancer survival correlated with OMA1 gene expression levels in multiple studies.^9, 58, 59^ The protooncogene MYC could thereby be responsible for OMA1’s differential expression in tumors.^9^ Another study placed OMA1 downstream of tumor protein p53.^55^ On the other hand, OMA1 and its substrate OPA1 are also connected to heart disease.^1–8^ Considering that kinase inhibitors are notorious for cardiotoxicity^60, 61^, one could speculate about cross-reactivity with the OMA1 protease.

## METHODS

**All methods are provided in the Supporting Information.**

## ACKNOWLEDGMENTS

Thank you for the generous support of my research by the National Institute on Aging (NIA) of the National Institutes of Health (NIH) under Award Number R43AG063642. I am indebted to the NIH’s National Cancer Institute (NCI) for sharing screening libraries and compound collections free of charge.

## CONFLICT OF INTEREST

Dr. Marcel V. Alavi is shareholder of 712 North Inc., a California-based pharmaceutical company.

