## Supporting Information for "OMA1 High-throughput Screen Reveals Protease Activation by Kinase Inhibitors"

Marcel V. Alavi

712 North Inc., Berkeley, CA

**Corresponding address:**

712 North Inc.

Berkeley CA-94720, USA

Electronic address:

### SUPPLEMENTARY FIGURES

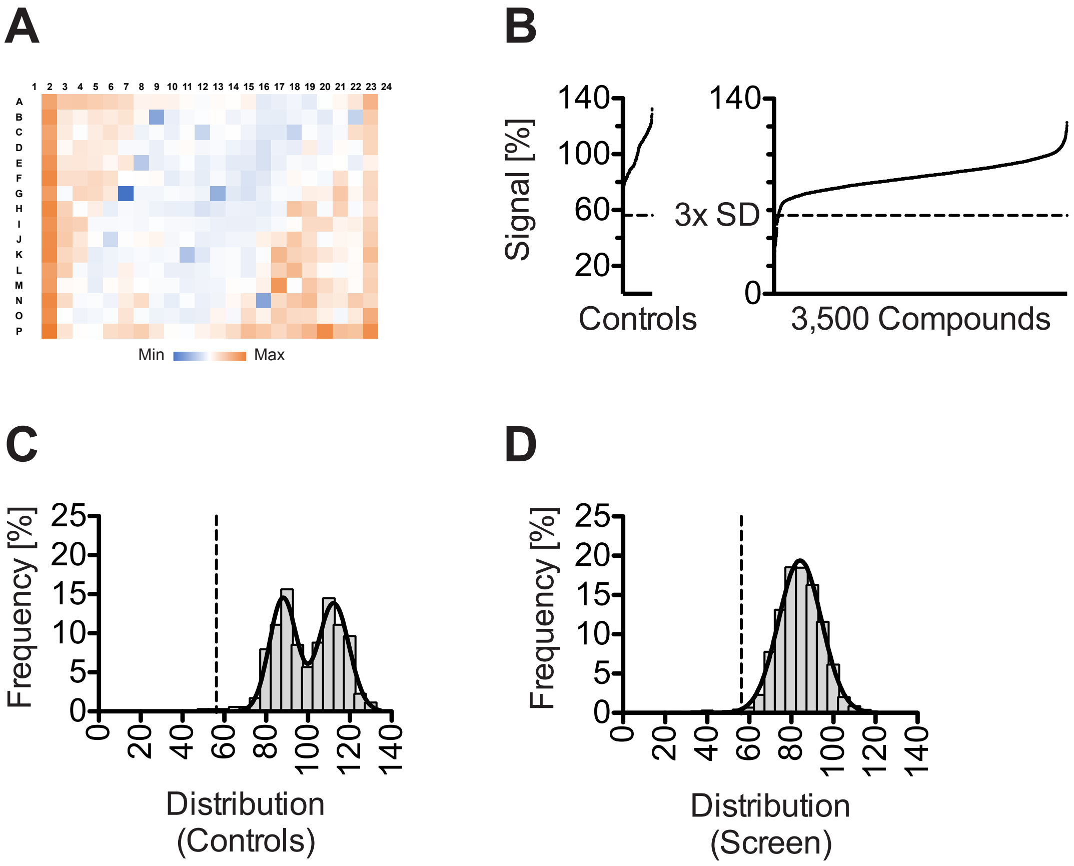


**Supplementary Figure S1: 3,500 molecules sampled with Luke cells to estimate assay interference.** Luke cells were exposed to 3,500 compounds in search of molecules that would lower luminescence. A heatmap illustrates the plate drift in Luke assays during the recording due to high substrate turnover of the luciferase (**A**). A signal reduction of at least 3 SD of untreated controls was considered significant (**B**). The 352 controls were distributed following the sum of two Gaussian distributions, which corresponded to the controls in column #2 and column #23, respectively (**C**; $R^2$=0.96). The signal of the 3,500 molecules were distributed normally with 56 molecules crossing the significance level (**D**; $R^2$=1.00).

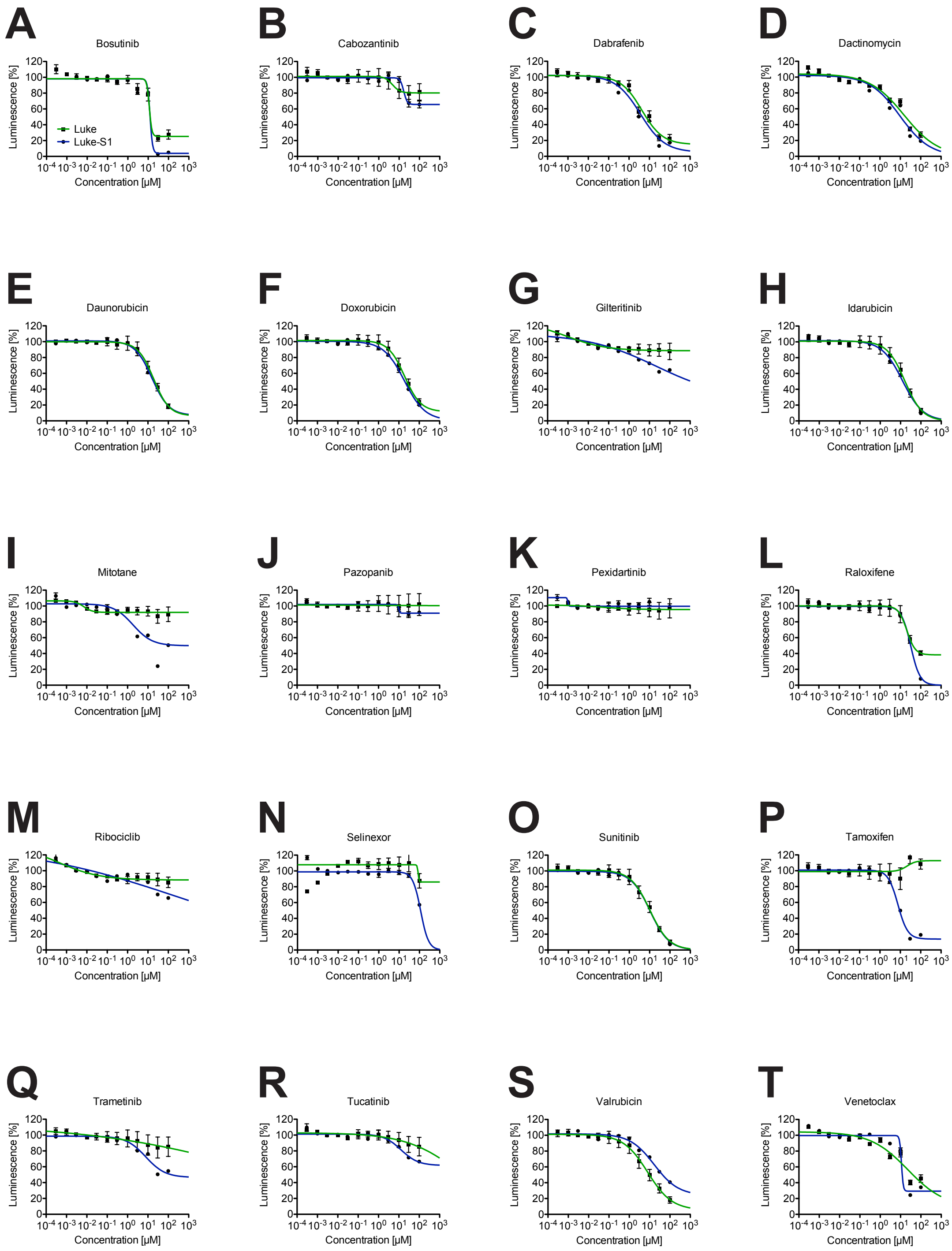


**Supplementary Figure S2: Follow up on 20 cancer drugs identified during the Luke-S1 screen.**

Dose-response curves for the denoted cancer drugs in Luke-S1 assays (blue line) and Luke control assays (green line). Note, that most of these drugs show assay interference.

### METHODS

#### Cell culture

Hek293T human embryonic kidney cells were obtained from the cell culture core facility at UC Berkeley. All cell culture reagents were from Gibco (Invitrogen). Cells were grown under standard conditions at 37°C in a humidified atmosphere with 5% $CO_2$ in advanced DMEM (#12491023) supplemented with 5% (v/v) fetal bovine serum (#10437028), 1× GlutaMAX (#35050079), and 100 units/ml of penicillin and 100 µg/ml streptomycin (#15140122).

#### Reporter cell lines

The DNA sequences of all reporter constructs were codon-optimized for human expression, synthesized and cloned into a mammalian expression vector under the control of a CMV promotor. 80% confluent Hek293T cells in a 12-well plate were transfected with 1 µg of plasmid and 3 µl of X-tremeGENE 360 (Roche Diagnostics) following the manufacturer's protocol. 24 hours post transfection cells were switched to selection media containing 2.5 µg/ml puromycin and from thereon cultured in the same. For the reporter assays, cells were expanded in T225 flaks, trypsinized and harvested by centrifugation at 200 × g for 2 minutes. Cells were resuspended in DMEM/F12 (#11039047, Gibco) without supplements and the volume was adjusted to $10^6$ cells per ml.

**Chemicals**

Carbonyl cyanide m-chlorophenyl hydrazine (CCCP) was purchased from Sigma-Aldrich and valinomycin from Cayman Chemical Company. Screening libraries were obtained from the National Cancer Institute (NCI) through the NCI Experimental Therapeutics (NExT) Program and from the NCI's Open Chemical Repository through the Developmental Therapeutics Program (DTP). All compounds were stored as 10 mM stock solutions in dimethyl sulfoxide (DMSO) at -20˚C. CCCP and valinomycin working dilutions were prepared fresh in DMEM/F12 on the day of the experiment. The final concentration of the screening compounds was 10 µM with 0.1% (v/v) DMSO. Unless stated otherwise, 100 nM valinomycin in DMEM/F12 for 30 minutes was used to activate OMA1 in all experiments.

**Gene silencing**

OMA1 protein knockdown was accomplished by silencing gene expression with pooled siRNA targeting human OMA1. 50% confluent 293T Luke-S1 cells (about 125,000 cells per well of a 24-well plate) were transfected with 0.8 µg OMA1 siRNA (#EHU072451, Sigma-Aldrich) or fluorescent universal negative control siRNA (#SIC007, Sigma-Aldrich) using X-tremeGENE 360 transfection reagent following the manufacturer's protocol. Cells were investigated 36 hours post transfection.

**Protein analysis**

Whole cell extracts were prepared by harvesting cells in lysis buffer A comprising 25 mM Tris-HCl (pH 7.6), 150 mM NaCl, 0.1% NP-40, 0.1% sodium deoxycholate, 0.01% SDS, 1× HALT protease inhibitor cocktail (ThermoFisher Scientific). Whole cell lysates were cleared from debris by 20,000 × g centrifugation for 5 minutes at 4˚C. The supernatant (about 30 μg total protein per lane) was separated under reducing conditions on tris-glycine mini-gels (Invitrogen) following the manufacturer's protocol, blotted onto PVDF membranes, blocked with 5% (w/v) dry milk powder in TBST (tris buffered saline with 0.1% Tween20), and labeled with the denoted antibodies diluted in blocking solution following standard procedures. The primary antibodies were: LgBiT (#N7100, Promega; 1:500), OMA1 (#SC-515788, Santa Cruz Biotechnology; 1:500), and OPA1 (#612606, BD Biosciences; 1:1,000). Alkaline-phosphatase-conjugated secondary antibodies (1:5,000) and 1-step NBT/BCIP substrate (ThermoFisher Scientific) were used for protein detection. Mitochondria-enriched fractions were prepared by breaking open cells on ice with a snuggly-fitting Potter-Elvehjem homogenizer in hyperosmotic buffer B (25 mM Tris-HCl (pH 7.6), 150 mM NaCl, 70 mM sorbitol, 5 mM EDTA, 1 mM DTT, 1× HALT protease inhibitor cocktail). Crude cell extracts were cleared from debris by 3-minute centrifugation at 1,000 × g. Mitochondria were sedimented by centrifugation at 15,000 × g for 30 minutes at 4˚C and then resuspended in buffer A. The supernatant and the mitochondria-enriched pellet of approximately 30 μg total cellular protein were analyzed by gel electrophoreses and Western-blotting as described above.

**Immunocytochemistry**

Cells were grown over night in chambered microscope slides under standard conditions and exposed to 1× MitoLite Red FX600 (Cayman Chemical Company) in culture media for another 30 minutes before they were fixed in ice-cold methanol for 10 minutes. Cells were washed twice in PBS and blocked for 45 minutes with 3% (w/v) BSA in PBS and incubated with the LgBiT antibody (1:50 in blocking solution) at 4˚C over night. Cells were labeled the next day for 1 hour with a goat α-mouse SureLight 488-conjugated secondary antibody (Columbia Biosciences; 1:250 in blocking solution) and mounted in ProLong Diamond Antifade with DAPI (ThermoFisher Scientific). Images were acquired with a Zeiss Axio Observer.A1 with brightness and contrast adjusted post acquisition.

**Cell assays**

For OMA1 activator screens, 20,000 reporter cells in 20 µl DMEM/F12 were dispensed with a µFill instrument (BioTek) into each well of a 384-well plate (#3704, Costar, Corning Inc.). The wells of columns #3 to #22 contained 5 µl of 50 µM test compound in 0.5% (v/v) DMSO in DMEM/F12 (10 µM final concentration). Wells of columns #2 and #23 served as controls and contained 5 µl DMEM/F12 only. Cancer drugs were tested in the same manner with the only difference that 20 µl cells were added to 1 µl of 200 µM drug in 2% (v/v) DMSO in water. The 384-well plates were incubated for 1 hour at 37˚C in a humidified $CO_2$ incubator. After the incubation step, 10 µl of 1:250 luciferase substrate (Nano-Glow #N113, Promega) in DMEM/F12 were added to each well with the build-in auto-dispenser of a Fluoroskan FL plate reader

(ThermoFisher Scientific). Plates were mixed, briefly spun down, and bioluminescence was measured with said device at 25˚C using an integration time of 500 milliseconds.

For the OMA1 inhibitor screens, cells were dispensed into 384-well plates with 5 µl 50 µM test compounds as described above and preincubation for 60 minutes at 37˚C in a humidified $CO_2$ incubator. After the preincubation step, 5 µl of 600 nM valinomycin (100 nM final concentration) were added to each well and incubated for an additional 30 to 60 minutes. Controls were left untreated. 10 µl of 1:250 luciferase substrate was then added and bioluminescence measured as described above.

For follow-up assays and the initial characterization of the 3 reporter cell lines, about 25,000 cells per well were seeded in 96-well half-area plates (#3688, Costar) and allowed to attach for at least 6 hours. Media was then carefully removed and the cells were exposed to the denoted experimental conditions in 30 µl volume. Bioluminescence was measured after adding 3 µl 1:100 luciferase substrate as described above.

**Data analysis**

Western blots were quantified by densitometry using NIH's imageJ software [1]. Statistical analyses were performed with GraphPad Prism 5.0c (GraphPad Software Inc.). All results are represented as mean ± standard deviation (SD). Groups from at least 3 independent experiments were compared by Student's t-test or using an analysis of variance (ANOVA). Differences between groups were considered to be significant at $p$ values of ≤0.05. Michaelis-Menten enzyme kinetics ($K_M$) were calculated from two independent substrate titrations assuming the proprietary substrate concentration is 250 µM. This assumption was based on two studies which estimated NanoLuc's $K_M$ at 10 µM and 1:25 (0.04×) substrate dilution, respectively [2, 3]. Dose-response relationships were plotted as semi-logarithmic graphs and half maximal effective concentrations ($EC_{50}$) were calculated using a non-linear inhibitor regression model with variable Hill-slope. The data from the different screens was normalized and displayed in a rank order and a frequency distribution histogram with an automatically determined bin width.
